# High Frequency Terahertz Stimulation Alleviates Neuropathic Pain by Inhibiting the Pyramidal Neuron Activity in the Anterior Cingulate Cortex of mice

**DOI:** 10.1101/2024.03.06.583763

**Authors:** Wenyu Peng, Pan Wang, Chaoyang Tan, Han Zhao, Kun Chen, Huaxing Si, Yuchen Tian, Anxin Lou, Zhi Zhu, Yifang Yuan, Kaijie Wu, Chao Chang, Yuanming Wu, Tao Chen

## Abstract

Neuropathic pain (NP) is caused by a lesion or disease of the somatosensory system and is characterized by abnormal hypersensitivity to stimuli and nociceptive responses to non-noxious stimuli, affecting approximately 7–10% of the general population. However, current first-line drugs like non-steroidal anti-inflammatory agents and opioids have limitations, including dose-limiting side effects, dependence, and tolerability issues. Therefore, developing new interventions for the management of NP is urgent. In this study, we discovered that the high-frequency terahertz stimulation (HFTS) at approximate 36 THz effectively alleviates NP symptoms in mice with spared nerve injury. Computational simulation suggests that the frequency resonates with the carbonyl group in the filter region of Kv1.2 channels, facilitating the translocation of potassium ions. *In vivo* and *in vitro* results demonstrate that HFTS reduces the excitability of pyramidal neurons in the anterior cingulate cortex through enhancing the voltage-gated K^+^ and also the leak K^+^ conductance. This research presents a novel optical intervention strategy with terahertz waves for the treatment of NP and holds promising application in other nervous system diseases.

## Introduction

Neuropathic pain (NP) refers to a debilitating chronic pain condition, which is often a consequence of nerve injury or of the diseases such as cancer, diabetes mellitus, infection, autoimmune disease, and trauma (*^1, 2^*). The symptoms of NP include spontaneous pain, hyperalgesia and mechanical allodynia. Unfortunately, NP is often resistant to currently available drug treatments, including non-steroidal anti-inflammatory drugs and even opioids (*^3^*). More evidences reveal that NP is not merely a symptom of a disease but rather an expression of pathological operations of the nervous system (*^4^*). Therefore, developing new therapeutic technology aimed at these underlying mechanisms for pain relief represents a considerable challenge.

Compared with the limitations of chemical-based drugs research, physics-based treatment offers a new concept and opportunity for intervening in NP. Optogenetics, as an interdisciplinary approach, has demonstrated therapeutic potential in NP. However, the limitations of viral vector delivery systems in humans are well-known (*^5^*). Recently, evidence has emerged suggesting that high frequency terahertz (THz) photons directly resonate with molecules, thereby regulating corresponding biological functions. For instance, our previous study demonstrated that a 34.88 THz wave resonates with Aβ protein, disrupting the process of fibril formation (*^6^*). Li et al. discover that the band of 42.55 THz resonates with the stretching mode of either the –COO- or the –C=O group significantly enhancing the Ca^2+^ conductance (*^7^*). Zhu et al. conclude that 48.2 THz photons greatly increase the permeability of sodium channel by a factor of 33.6 through breaking the hydrated hydrogen bonding network between the hydrosphere layer of the ions and the carboxylate groups (*^8^*). Additionally, the frequency of 53.5 THz has been reported to enhance the voltage-gated K^+^ currents, which modulate the startle response and associative learning (*^9, 10^*). These studies strongly prompt us the potential application of THz photons in the treatment of neuropathic pain by targeting the ion channels (*^11^*).

The anterior cingulate cortex plays a crucial role in pain regulation (*^12^*). Our previous research has demonstrated that nociceptive information resulting from nerve injury is transmitted to the ACC (*^13^*). This region exhibits pre- and postsynaptic long-term plasticity (LTP), which contributes to chronic pain and associated negative emotions (*^14, 15^*). Furthermore, descending projection pathways from the ACC enhance the neuronal activity of the spinal dorsal horn (SDH) and regulate nociceptive sensory transmission (*^16^*). Brain-imaging and MRI studies also provide evidence of hyperexcitability in the ACC during both acute and chronic pain (*^17, 18^*). Specifically, the activity of pyramidal cells in the ACC is directly correlated with the expression of chronic pain (*^19, 20^*). Optogenetic excitation of ACC pyramidal cells induces pain, while their inhibition leads to analgesia (*^21^*). Therefore, targeting the cortical regions of the ACC and inhibiting the activity of ACC pyramidal neurons may hold promise as a strategy for treating NP (*^22–25^*).

Neuronal excitability is influenced by various types of voltage-gated ion channels and among them, voltage-dependent potassium (Kv) channels, as one of the important physiological regulators of neuronal membrane potentials, has been proposed as potential target candidates for pain therapy (*^26–28^*). Zhao et al. have reported that enhancing Kv currents in injured dorsal root ganglion (DRG) neurons alleviates neuropathic pain (*^29^*). Additionally, Fan et al. have demonstrated that lumbar (L)_5_ spinal nerve ligation (SNL) leads to a time-dependent decrease in Kv1.2-positive neurons in the ipsilateral L_5_ DRG. However, rescuing Kv1.2 expression in the injured L5 DRG attenuates the development and persistence of pain hypersensitivity (*^30^*). These findings highlight the potential of targeting Kv channels as a therapeutic approach for managing pain.

In this study, we investigated the effects of high-frequency terahertz stimulation (HFTS) on the Kv model. By analyzing the absorbance spectra of Kv1.2 channels, we observed a significant response to photons with a frequency of approximately 36 THz. This frequency modulates the resonance of the carbonyl group in the Kv1.2 structure, affecting the action potential waveform and frequency as demonstrated through simulations. Subsequently, we conducted *in vivo* multi-channel recordings and *in vitro* patch recordings to confirm the activation effect of HFTS on K^+^ conductance and its inhibition of neuronal activity in the ACC pyramidal cells. Importantly, the application of HFTS resulted in a significant reduction in pain behavior in mice with spared nerve injury (SNI).

## Results

### HFTS attenuates the generation of action potential through molecular dynamics simulation

To identify a specific terahertz (THz) frequency capable of modulating a major subset of voltage-gated potassium (Kv) channels, we developed an integrated model comprising both mouse Kv channels (Protein Data Bank [PDB] ID: 3LUT) and Na^+^ channels (PDB ID: 3RVY) (Fig. 1a and Supplement Fig. S1). We conducted a comprehensive analysis of the spectral absorption characteristics within the THz frequency range. Our results revealed a pronounced absorption peak at approximately 36 THz for the potassium channel, which exhibited a considerable board band compared to the absence of a corresponding peak for the sodium channel (Fig. 1a). This indicates a preferential and resonant absorption of photons at the ∼36 THz frequency by the potassium channel. We then tested the possible kinetic changes of Kv1.2, the typical and widely distributed Kv channel in the central nervous system (*^31^*), following the absorption of these THz photons. Our findings indicated a significant kinetic change in the -C=O groups of the channel filter structure, as evidenced by an expansion of its van der Waals radius by approximately 0.5 Å (Fig. 1b). Furthermore, during exposure to THz photons, the conductance of the potassium ion channel exhibited an almost linear increase with the intensity of the THz field, while the conductance of sodium ions remained largely unchanged (Fig. 1c). Interestingly, under terahertz photonic influence, the cortical neurons model showed significantly decreased in discharge (Fig. 1d). We performed a detailed waveform analysis of the action potentials, including parameters such as full width at half maximum (FWHM) and firing frequency (Fig. 1e). Our observations revealed that the FWHM of action potentials in the THz-exposed group decreased to 95% of the control group (Fig. 1f, red column), and the firing frequency experienced a reduction of approximately 70% after THz photon stimulation (Fig. 1f, green column). These results collectively suggest that THz photons primarily attenuate neuronal firing activity by increasing potassium ion conductance, thereby modulating neuronal excitability.

**Fig. 1.**
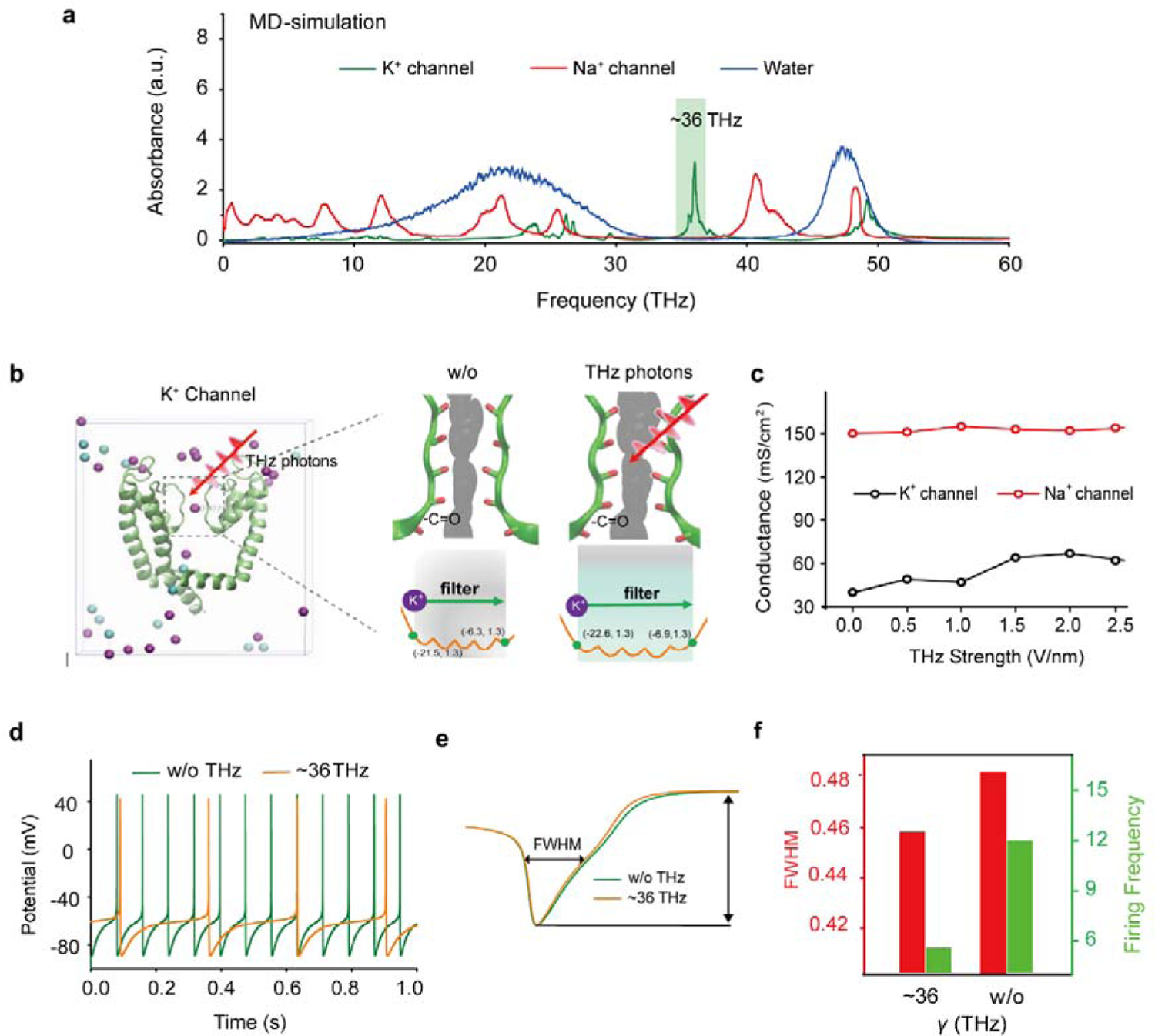
Specific frequency THz photons resonate K_v_ channel and decrease the AP firing rate in cortical neurons through molecular dynamics simulation. (a) Absorbance spectra of voltage-gated potassium/sodium ion channels and the bulk water. (b) The dynamic attributes of the Kv1.2 filter structure in pre- and post-exposure to HFTS. Purple balls represent the K^+^, blue balls represent the Cl^-^. (c) The alterations in potassium/sodium ion conductance consequent to the influence of HFTS. (d) Changes of the firing rate of APs of cortical neuron models before and after HFTS. (e) The FWHM of an AP pre- and post HFTS. (f) Changes in FWHM and firing frequency with or without HFTS. HFTS, high frequency terahertz stimulation. AP, action potential. FWHM, Full Wide of Half Maximum.

### HFTS enhances voltage-gated K^+^ currents and leak K^+^ currents of pyramidal neurons in the ACC

To investigate the impact of high-frequency terahertz stimulation (HFTS) on voltage-gated potassium/sodium (Kv/Nav) channels, which play a crucial role in action potential generation and waveform, we conducted whole-cell patch recording from layer-5 pyramidal neurons (PYR^ACC^) in acute slices of anterior cingulate cortex (ACC) in mice with spared nerve injury (SNI) (Fig. 2a). Initially, we examined the Nav current by applying a series of test pulses (from −80 to −10 mV) with a command voltage of −100 mV (20 ms) (Fig. 2b). Upon illumination with ∼36 THz photons (0.3 ± 0.05 mW) for durations of 5, 10 and 20 minutes, we observed that HFTS had no significant effect on the activation and inactivation curve slope, half-activation and half-inactivation voltage and time constants (tau) for half-activation and half-inactivation voltage (Figs. 2c-f). These findings indicate that HFTS does not affect Nav channel-mediated currents. Subsequently, we investigated the influence of HFTS on Kv currents by applying a series of test pulses (100 ms) ranging from −70 to +130 mV with a command voltage of −100 mV (Fig. 2g). Our results demonstrated that the application of HFTS induced a significant increase in the amplitude of K^+^ currents (Table S1-1) and an enhanced slope of the current-voltage characteristic (I-V) curve (Figs. 2h-j), without affecting the half-activation voltage (Fig. 2k). Furthermore, to assess the duration of neuronal effects induced by HFTS (15 minutes), we examined Kv currents at 5 minutes and 20 minutes post-HFTS. It was observed that the Kv current were enhanced 20 minutes post-HFTS (Fig. 2l, Table S1-2). Simultaneously, we studied the effect of HFTS on K_Leak_ currents by applying a series of test pulses (400 ms) ranging from −120 to −30 mV followed with a command voltage of −70 mV (Fig. 2m). Currents recorded at and above −65 mV membrane potential were analyzed for potential K_Leak_ currents (*^32^*). We found that HFTS also induced a significant increase of K_Leak_ currents at holding potential at −40 and −30 mV (Fig. 2n) (Table S1-3). These experiments demonstrated that HFTS at approximately 36 THz not only influenced Kv currents but also affected K_Leak_ channel activity, resulting in an acceleration of potassium ion flow and an increase in potassium conductance in PYR^ACC^ neurons. Importantly, these experimental findings were consistent with the results obtained from molecular dynamics analysis.

**Fig. 2.**
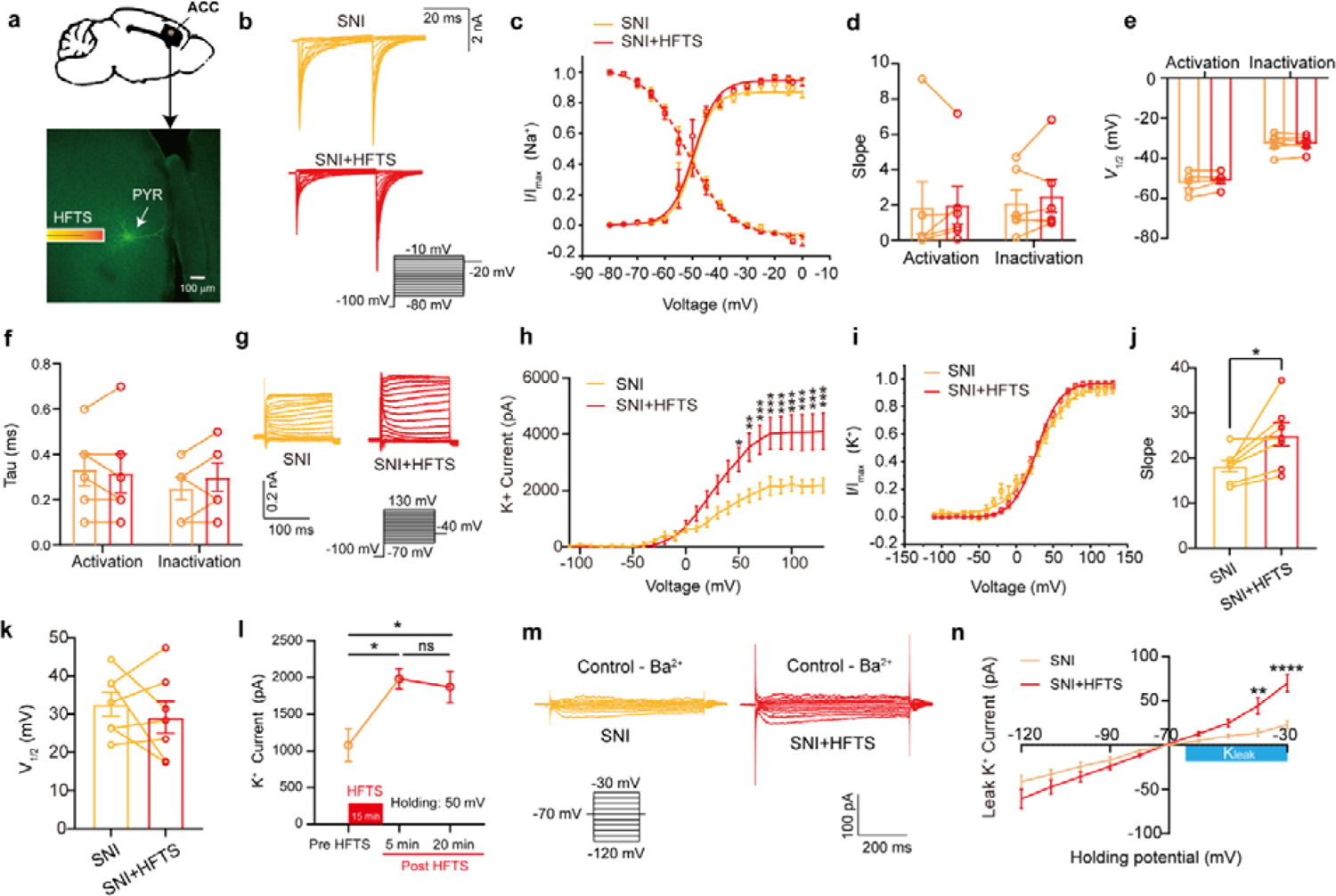
HFTS enhances Kv and K_Leak_ currents of pyramidal neurons in SNI mouse *in vitro*. (a) Anatomical location of ACC region in mice and a recorded PYR neuron (biocytin-labeled, green). (b) Representative Nav currents without (orange) or with HFTS (red) under the given step voltage protocol. (c) The activation and inactivation curves of Nav currents with and without HFTS. (d-f) The corresponding slopes of the activation and inactivation curves (d), the comparison of the half-activation and inactivation voltages (e) and the time constants (tau) of half-activation voltage/half-inactivation voltage (f). (g) Representative Kv currents evoked by a series of step voltages (inset) without (orange) or with HFTS (red). (h) I-V plots constructed from the values of traces shown in (g). (SNI *vs.* SNI + HFTS: *F_(1, 10)_* = 6.846, *P* < 0.0001, n_SNI_ = 6, n_SNI+HFTS_ = 6; Two-way ANOVA followed by *post hoc* comparison using the Šídák’s multiple comparisons test). (i) The activation curves of the Kv currents with and without HFTS. (j) The corresponding slopes of the activation curves (SNI *vs*. SNI + HFTS: *t* = 5.872, *P* = 0.0011, n = 7, unpaired *t*-test. **, *P* < 0.01, ****, *P* < 0.0001). (k) the half-activation voltages of the activation curves. (l) Changes in the impact of Kv current post-HFTS. (*F_(4, 15)_* = 4.19, *P* = 0.0178, n = 4; One-way ANOVA followed by *post hoc* comparison using the Šídák’s multiple comparisons test). (m) Representative K_Leak_ currents evoked by a series of step voltages (inset) without (orange) or with HFTS (red). (n) I-V plots constructed from the values of traces shown in (m). (SNI *vs.* SNI + HFTS: *F_(1, 12)_* = 1.688, *P* = 0.2182, n_SNI_ = 7, n_SNI+HFTS_ = 7; Two-way ANOVA followed by *post hoc* comparison using the Šídák’s multiple comparisons test).

### HFTS reduces the spike frequency of pyramidal neurons in the ACC

We proceeded to investigate the impact of the specific resonant frequency of THz photons on the excitability of PYR^ACC^ neurons in SNI and sham mice. Using whole-cell current-clamp recording, we compared the input-output curves of evoked action potentials before and after HFTS. Our findings revealed a significant increase in the spike frequency in SNI mice (Fig. 3a) (Table S2-1), which was effectively rescued by the application of HFTS (Fig. 3b) (Table S2-2), but not by 465 nm blue light stimulation (BLS) (Fig. 3c) (Table S2-3). Furthermore, the spike frequency in sham mice also decreased after the application of HFTS (Fig. 3d) (Tables S2-4). To further analyze the properties of single action potentials, we induced them by applying a depolarizing current pulse (30 ms) of an appropriate suprathreshold magnitude (Fig. 3e). In SNI mice, we observed a decrease in the rheobase and an elevation in the resting membrane potential (RMP) compared to those in sham mice. However, these alterations were reversed by the application of HFTS, while BLS had no effects. Other parameters, such as voltage threshold, amplitude and half-width of the action potentials, were not different between SNI or sham mice with and without HFTS (Figs. 3f-k). Given that the spike firing, rheobase and RMP are closely related to low-threshold Kv channels and K_Leak_ channels (*^26, 31, 33^*), these results suggest that HFTS affects the activity of PYR^ACC^ neurons through its specific impact on Kv and K_Leak_ channels.

**Fig. 3.**
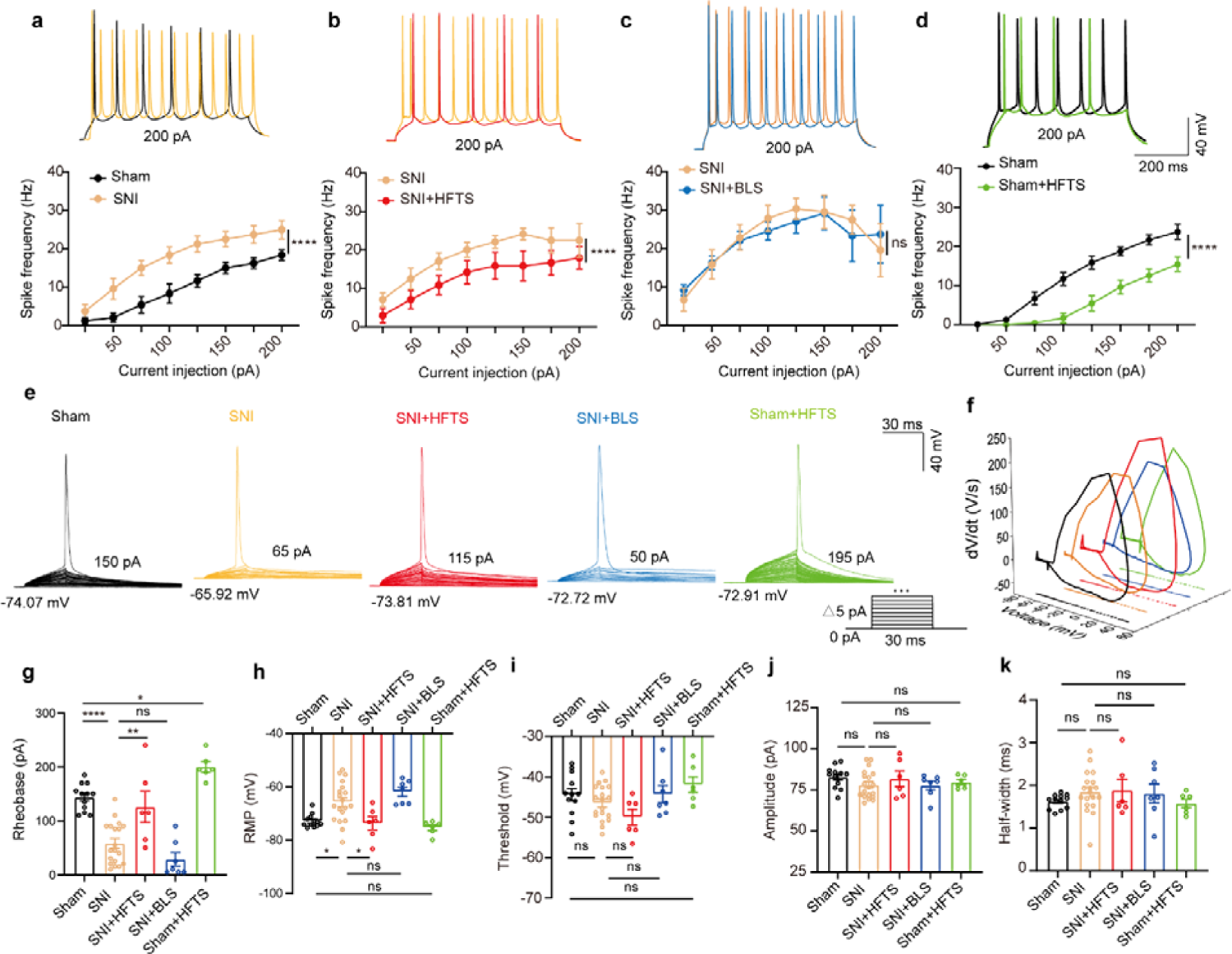
HFTS reduces the APs firing rate of pyramidal neurons in SNI and sham mice *in vitro*. (a-d) Representative traces (upper panels) and line-charts (lower panels) showing the changes of evoked spikes of pyramidal neurons in different groups. (Sham *vs*. SNI: *F_(1, 40)_* = 124.2, *P* < 0.001, n_sham_= 6, n_SNI_=6; SNI *vs.* SNI + HFTS: *F_(1, 40)_* = 23.13, *P* < 0.0001, n_SNI_=6, n_SNI+HFTS_=6; SNI *vs.* SNI + BLS: *F_(1, 40)_* = 0.1401, *P* = 0.7101, n_SNI_ = 6, n_SNI+BLS_ = 6; Sham *vs.* Sham + HFTS: *F_(1, 40)_* = 87.29, *P* < 0.0001, n_Sham_ = 6, n_Sham+HFTS_ = 6. Two-way ANOVA followed by *post hoc* comparison using the Šídák’s multiple comparisons test). (e) Superimposed traces showing the single AP evoked by threshold current stimulation in different groups. (f) Phase plots of AP traces in each groups. (g) Histograms showing the statistical comparison of rheobase in each group. (Sham *vs*. SNI: q = 8.456, *P* < 0.0001, n_sham_ = 12, n_SNI_ = 19; SNI *vs.* SNI + HFTS: q = 5.264, *P* < 0.01, n_SNI_ = 19, n_SNI+HFTS_ = 6; Sham *vs.* Sham + HFTS: q = 4.098, *P* < 0.05, n_SNI_ = 19, n_SNI+HFTS_ = 6. one-way ANOVA followed by *post hoc* comparison using the Tukey’s multiple comparisons test). (h) The RMP in each group (Sham *vs.* SNI: q = 4.887, *P* < 0.05, n_sham_ = 12, n_SNI_ = 19; SNI *vs.* SNI + HFTS: q = 4.29, *P* < 0.05, n_SNI_ = 19, n_SNI+HFTS_= 6; Sham *vs.* Sham + HFTS: q = 1.261, *P* > 0.05, n_SNI_ = 19, n_SNI+HFTS_ = 6. one-way ANOVA followed by *post hoc* comparison using the Tukey’s multiple comparisons test). (i-k) HFTS has no significant effect on the threshold, amplitude and half-width of APs in pyramidal neurons.*, *P* < 0.05, **, *P* < 0.01, ***, *P* < 0.001, ****, *P* < 0.0001, ns, *P* > 0.05. BLS, blue light stimulation.

### HFTS decreases the excitability of pyramidal neurons in the ACC *in vivo*

We then investigate the effect of HFTS on the activities of PYR^ACC^ in head-fixed awake SNI mice. One-week prior to the illumination experiment, a 16-channel electrode was implanted into the ACC (the detailed structure of this device shows in Fig. 4a). Then we applied THz photon stimulation for 15 minutes and compared the neuronal activities before and after HFTS (Fig. 4b). Our findings revealed a significant decrease in the mean firing rate of ACC neurons after HFTS application in both the sham and SNI groups (Figs. 4c and g). To further analyze the effect of HFTS on the PYR^ACC^, we classified them along with interneurons in the ACC (INT^ACC^) based on their firing rate, trough-to-peak duration and half width (Fig. 4d), as described in our previous study (*^19^*). We assessed the internal-spiking interval (ISI) and waveform characteristics of the isolated neurons in each channel to ensure that the pre- and post-HFTS units originated from the same neuron (Fig. 4e). In the sham group, we observed that 63.4% of PYR^ACC^ neurons exhibited a decrease in firing rate, 10.8% of PYR^ACC^ showed an increase, and 25.8% of PYR^ACC^ remained unchanged (93 well-isolated PYR^ACC^ neurons were recognized out of 108 total recorded units). In the SNI group, we found that 61.8% of PYR^ACC^ neurons exhibited decreased activity, 20.3% of PYR^ACC^ showed increased activity, and 17.9% of PYR^ACC^ remained unchanged (123 well-isolated PYR neurons were recognized out of 130 total recorded units) (Fig. 4f). Consistently, the increased mean firing rate of PYR^ACC^ neurons in SNI mice was significantly inhibited by the application of HFTS (Fig. 4h). The activity of INT^ACC^ also tended to decrease after HFTS (Fig. 4i). In contrast, blue light stimulation (BLS) has no effect on the mean firing rate on the PYR^ACC^ and INT^ACC^ in both sham and SNI mice (Supplement Fig. S2). These results indicate that HFTS reduces the spike firing of ACC neurons, whereas BLS does not have the same effect.

**Fig. 4.**
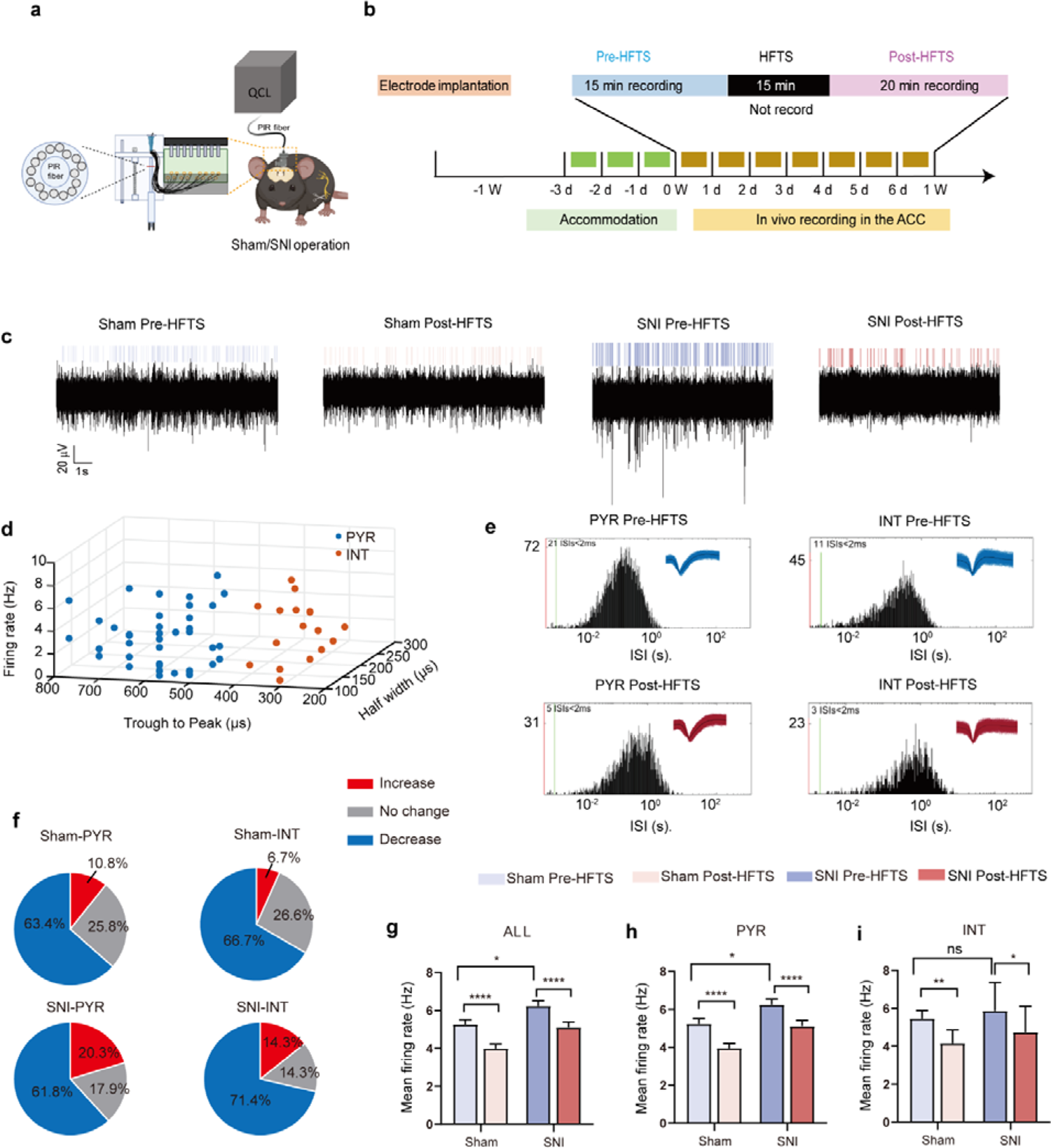
HFTS decreases the mean firing rate of pyramidal neurons in the ACC in both sham and SNI awake mice. (a) Schematic diagram of the single-unit recording of the ACC using an *in vivo* multi-channel recording technique. (b) The timeline and the stimulating pattern of HFTS on an awake mouse. (c) Example recording signals of ACC neurons before and after HFTS application in sham and SNI groups, respectively. (d) ACC neurons are classified as pyramidal (PYR) cells and interneurons (INT) using *k*-means cluster-separation algorithm based on their electrophysiological properties. (e) Histograms of the inter-spike intervals (ISI) from the spikes of a PYR and an INT in pre- and post-HFTS recording period. Insets at the top right corner show the waveforms of the detected single unit. (f) Pie charts summarize the changes in firing rate of PYR and INT in sham and SNI groups. Pre *vs.* post HFTS, Wilcoxon rank-sum test. (g) The mean firing rate of all recorded neurons in sham and SNI groups before and after HFTS. Sham group (*P* < 0.0001, n = 108, Wilcoxon matched-paired signed rank test), SNI group (*P* < 0.0001, n = 130, Wilcoxon matched-paired signed rank test), SNI pre-HFTS vs. Sham pre-HFTS (*P* = 0.0447, Mann-Whitney test). (h) The mean firing rate of PYR neurons in sham and SNI groups before and after HFTS. Sham group (*P* < 0.0001, n = 93, Wilcoxon matched-paired signed rank test), SNI group (*P* < 0.0001, n = 123, Wilcoxon matched-paired signed rank test), SNI pre-HFTS vs. Sham pre-HFTS (*P* = 0.0274, Mann-Whitney test). (i) The mean firing rate of INT neurons in sham and SNI groups before and after HFTS. Sham group (*P* = 0.0084, n = 15, Wilcoxon matched-paired signed rank test), SNI group (*P* = 0.0313, n = 7, Wilcoxon matched-paired signed rank test), SNI pre-HFTS vs. Sham pre-HFTS (*P* = 0.3322, Mann-Whitney test). *, *P* < 0.05, **, *P* < 0.01, ****, *P* < 0.0001, ns, *P* > 0.05.

### HFTS alleviates mechanical allodynia of SNI mice

Finally, we tested whether application of HFTS into the ACC induced analgesic effects. The SNI surgery and optic fiber tube implantation into the ACC were performed one week before pain behavioral tests, which included the mechanical pain threshold test and Catwalk analysis (Fig. 5a). We compared the paw withdrawal mechanical thresholds (PWMTs) before and after HFTS (0.3 ± 0.05 mW at the tip of the optic fiber) application for 15 minutes and found that SNI treatment significantly decreased the PWMTs compared to the sham group. However, after the application of HFTS, the PWMTs significantly increased, even surpassing those in the sham group in the first 30 minutes (Fig. 5b). The analgesic effect lasted for around 160 minutes with a 15-minute application of HTFS and lasted for around 140 minutes with a 10-minute application of HTFS, suggesting a correlation between the duration of analgesia and the intensity of stimulation (Fig. 5c). In contrast, the PWMTs did not significantly change in the SNI group with the application of 465 nm blue light.

**Fig. 5.**
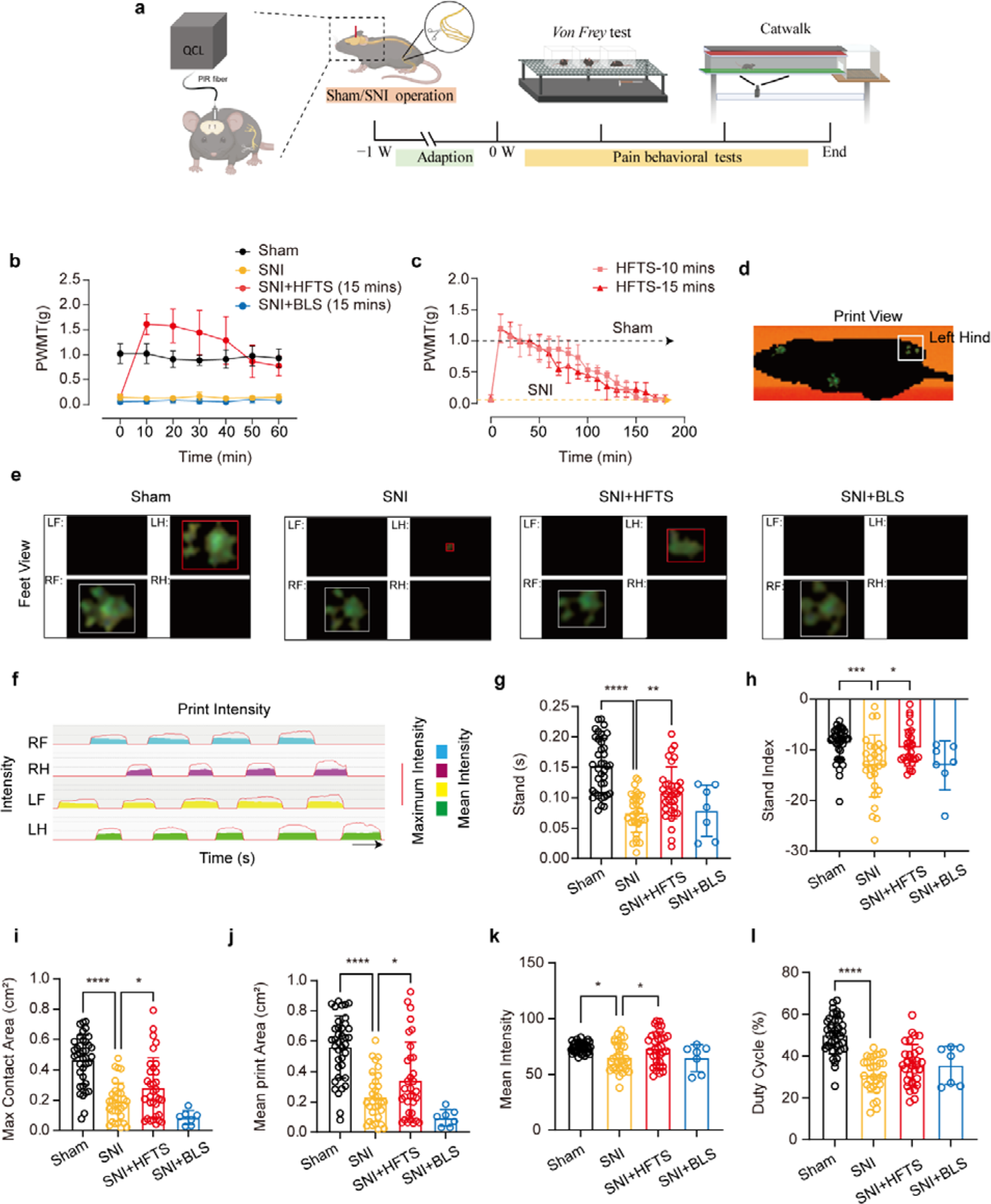
HFTS alleviates neuropathic pain of SNI mice through pain behavior tests. (a) Schematic of the establishment of NP model, the application of HFTS in ACC region and the following behavior tests including *Von Frey* test and Catwalk analysis. (b) HFTS increases the paw withdrawal mechanical thresholds (PWMTs) compared to the SNI model (*F_(18, 140)_* = 12.65. *P* < 0.0001. Sham *vs.* SNI: *P* < 0.0001; SNI *vs.* SNI + HFTS: *P* < 0.0001; n = 6 in each group. Two-way ANOVA repeated measures followed by *post hoc* comparison using the Šídák’s multiple comparisons test). (c) Duration of the analgesic effect with HFTS for 10 mins and 15 mins. (d) The print view of a mouse. (e) The feet view of the left front (LF), left hind (LH), right front (RF) and right hind (RH) in the groups of sham, SNI, SNI + HFTS and SNI + BLS, respectively. (f) The step sequence of a sham mouse who passing through the glass pane, the red line represents the maximum intensity of each foot, the color box represents the mean intensity of the corresponding print during walking. (g) HFTS increases the LH stand time of SNI mice (sham *vs.* SNI: *P* < 0.0001; SNI *vs.* SNI + HFTS: *P* < 0.01). (h) HFTS increases the LH stand index of SNI mice (sham *vs.* SNI: *P* < 0.001; SNI *vs*. SNI + HFTS: *P* < 0.05). (i) HFTS increases the LH max contact area of SNI mice (sham *vs.* SNI: *P* < 0.0001; SNI *vs*. SNI + HFTS: *P* < 0.05). (j) HFTS increases the LH mean print area of SNI mice (sham *vs.* SNI: *P* < 0.0001; SNI *vs*. SNI + HFTS: *P* < 0.05; SNI vs. SNI + BLS: *P* < 0.05). (k) HFTS increases the LH mean intensity of SNI mice (sham *vs.* SNI: *P* < 0.05; SNI *vs*. SNI + HFTS: *P* < 0.05). (l) HFTS have no significant for the pain behavior parameter of the duty cycle. *, *P* < 0.05, **, *P* < 0.01, ***, *P* < 0.001, ****, *P* < 0.0001. One-way ANOVA (f-k) followed by *post hoc* comparison using the Tukey’s multiple comparisons test. n_Sham_ = 38, n_SNI_ = 35, n_SNI+HFTS_= 34, n_SNI+BLS_ = 9.

Furthermore, we performed the Catwalk gait analysis (Figs. 5d-l), which provides exquisite and reliable observations for evaluating the spontaneous pain behaviors (*^34^*). We focused on the print intensity and print area related parameters of the left hind paw (ipsilateral side of the injured nerve). We found that SNI treatment significantly altered the stand time, the stand index, the max contact area, the mean print area, the mean intensity and the duty cycle (Figs. 5g-l). This suggests that the SNI mice tend to avoid standing and walking on their injured hind paw due to pain hyper-sensitivity. The application of HFTS but not BLS rescued most of the above parameters, indicating HFTS’ strong analgesic effect.

## Discussion

In the present study, we provide evidence that high frequency terahertz photons alleviate neuropathic pain in the SNI mice by decreasing the excitability of pyramidal neurons in the ACC. The mechanism underlying this effect is that HFTS increases voltage-gated potassium ion conductance through resonance with the carbonyl group in the potassium channel filter region (Fig. 6). Unlike optogenetic technology, HFTS can directly regulate the conformation of ion channel without delivering a transgene that encodes a light-response protein. It exhibits frequency selectivity and dependence to channel structure. This research suggests that HFTS has potential to serve as a novel optical technology for the treatment of NP pathology.

**Fig. 6.**
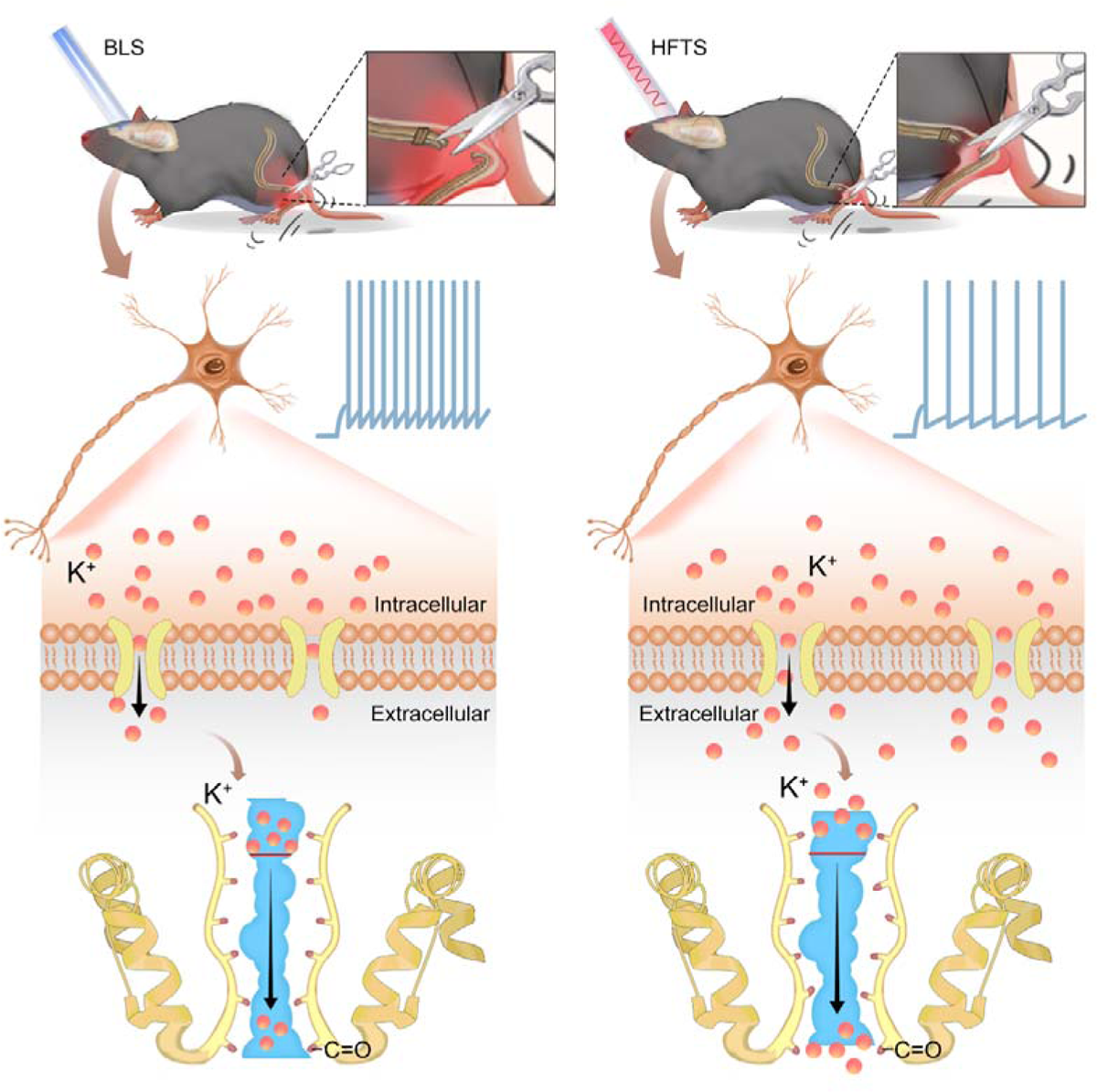
Schematic diagram shows the mechanism of HFTS in alleviating neuropathic pain. Left panel shows the group with BLS and the right panel shows the group with HFTS.

Neuropathic pain is closely associated with nociceptor excitability in the ACC (*^18, 35-37^*), with ion channels playing a fundamental role in determining neuronal excitability, particularly in the hyperexcitability of pyramidal neurons (*^11^*). Excitatory Nav channels, responsible for initiating and depolarizing the action potential, can be targeted by inhibitors to effectively decrease or eliminate electrical excitability. These inhibitors are commonly used in neurology as antiepileptic drugs. On the other hand, inhibitory K channels, responsible for repolarization, contribute to the initiation of action potentials in diverse ways (*^33^*). Enhancing K conductivity could have a similar effect to Nav channel blockers. For instance, retigabine, an activator for Kv7, has recently been approved as a first-in-class antiepileptic drug (*^38, 39^*). Among the 12 subfamilies of Kv channels (Kv1-12), Kv1.2 is the most prevalent in neuronal membranes(*^40^*) and has been reported to be significant associated with neuropathic pain (*^30, 41, 42^*). Although the response frequency of Kv1.2 at ∼53 THz (*^10^*) or ∼34 THz (*^43^*) and the corresponding modulation function have been verified due to the broad absorption band, this study highlights the significant resonance of Kv1.2 filter structure with photons at 36 THz. By optically stimulating ACC neurons with the frequency of ∼36 THz, we observed a significant reduction in the firing rate of pyramidal neurons’ action potentials in SNI mice, accompanied by a notable enhancement of K^+^ conductance. This confirms the effect of THz photons on Kv channels, including Kv1.2. However, to consolidate this conclusion, a more specific pharmacological experiment is necessary. For example, applying a blocking peptide to eliminate the Kv1.2 current and then testing whether this blocks the effects of HTFS would be a valuable test to perform in the future. Moreover, the application of HFTS resulted in significant changes in the rheobase and RMP of pyramidal cells, suggesting that HFTS may also affect the two-pore K^+^ channels (*^31^*). We thus tested the effect of HTFS on the K_leak_ current and confirmed this hypothesis. Additionally, it has been reported that basal excitability is influenced by the opening of low-threshold Kv1.2 channels, which filter out small depolarizations and thus control the number of triggered APs (*^40^*). However, we cannot exclude the possibility that THz photons affect other K channel functions, but further research is required to confirm this in the future.

During our research, we focus on studying of pyramidal neurons in the ACC (*^44^*). It has been reported that the firing rate of glutamatergic pyramidal cells, rather than inhibitory interneurons, increases in the ACC after chronic pain, suggesting an imbalance of excitatory/inhibitory (E/I) ratio (*^19, 35^*). In the local circuits of the ACC, inhibitory neurons release GABA and inhibit the activities of pyramidal cells. Different studies by Kang et al.(*^45^*) and Joseph Cichon et al.(*^46^*) have reported that specific activation of interneurons in the ACC or in the somatosensory cortex reduces pyramidal neuron hyperactivity and alleviate mechanical allodynia. Thus, the application of THz may induce complicated results through affecting the Kv channels distributed on both pyramidal cells and interneurons. However, as shown in our *in vivo* recording data (Fig. 4i), although THz illumination slightly decreased the activity of interneurons, which could potentially lead to an enhanced activity of local pyramidal cells, the direct and significant decrease in pyramidal cell activity caused by the illumination would overcome this disinhibitory effect, ultimately resulting in a net decrease in pyramidal cell activity. The behavioral analgesic effect caused by THz illumination also confirmed this conclusion.

There are several limitations in this study that should be acknowledged. Firstly, we did not investigate the thermal effect of HFTS on the Kv channel. Previous research has demonstrated the non-thermal, long-distance stimulation of high-frequency terahertz stimulation on neuronal activity (*^10^*). Nevertheless, the specific interaction between ∼36 THz and Kv channels in terms of thermal effects remains unexplored. Additionally, in this study, we used blue light as a comparison, and found no significant changes in potassium current and the excitability of pyramidal cells. This finding suggests the specificity of the terahertz frequency and supports the existence of non-thermal effects. Another limitation of our research is the use of an optic fiber to deliver the HFTS into the ACC region. This invasive approach may pose challenges for potential noninvasive applications. However, we believe that with the advancement of terahertz enhancement techniques, such as the use of metasurfaces or nanomaterials (*^47, 48^*), high-frequency terahertz waves show promising potential for broad applications in regulating diverse brain diseases, such as episodic ataxia (*^38, 49^*), benign familial neonatal convulsions (*^50^*), Alzheimer’s disease (*^51^*), and others.

## Material and methods

### Animals

Male adult (8–10 weeks) C57BL/6 were used for all experiments. Mice were housed on a 12 h light–dark cycle with food and water freely available. The living conditions were carefully controlled, with temperatures maintained at 22-26℃ and humidity at 40%. All animal procedures in the present experiments were in accordance with protocols approved by the Animal Care Committee of the Fourth Military Medical University. All efforts were made to minimize animal suffering and the number of animals used.

### Neuropathic Pain Model

We used spared nerve injury (SNI) model to establish neuropathic pain. The detailed process has been described in our previous study (*^19^*). In brief, mice were generally anaesthetized by 2% isoflurane. Three terminal branches of the left sciatic nerve were exposed by making a direct incision in the skin and a section of the biceps femoris muscle in the left thigh. The tibial nerve and the common peroneal nerve were ligated using 6-0 silk sutures and then sectioned distal to the ligation. After ligating and cutting the nerves, they were carefully put back into their original positions, and the muscle and skin were sutured in two layers. For the sham mice, animals only received an operation that exposed the branches of the left sciatic nerve but without any nerve injury. Following a week accommodation period, pain behaviors were assessed using the *von Frey* filament test and CatWalk gait analysis to confirm the successful establishment of the NP model.

### Molecular Dynamics Simulation

The simulation was conducted to gain a deeper understanding of the interaction between terahertz photons and ion channels. A composite model of mouse eukaryotic voltage-gated K^+^ channels (PDB ID: 3LUT) and eukaryotic Na^+^ channels (PDB ID: 3RVY) was built using the Charmm-GUI website. The model consisted of intact proteins, phospholipid bilayers, and saline solution (with a concentration of 0.15 M). Kinetic calculations were performed using GROMACS 5.1.2 software. The CHARMM 36 force field and periodic boundary conditions were applied to the proteins. Electrostatic interactions were handled using the connected element algorithm Ewald. During the simulation, the Rattle algorithm was used to constrain key lengths. The motion equation was solved using the Velocity-Verlet algorithm with a time step of 2 fs. Initially, the simulation was carried out at a room temperature of 303.5 K to observe the ion transport process within the channels at the molecular level. Subsequently, conductivity values (gNa, gK) for potassium and sodium ions and their corresponding absorption spectrum were calculated. To investigate the effect of ion transport under the influence of terahertz radiation, time-varying electric fields of THz radiation were added to the system. In the interaction of terahertz radiation with biological systems, electrical components play a crucial role. The electric field was used to simulate terahertz radiation, and its formula is as follows:

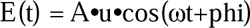

Where A represents the terahertz radiation intensity, u and phi represent the polarization direction and phase of the radiated photon, which set to (0, 0, 1) and 0, respectively. The terahertz radiation frequency v is related to the angular frequency ω by the equation:

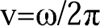

The cortical neuron Hodgkin-Huxley (H-H) model links the microscopic level of ion channels to the macroscopic level of currents and action potentials. The model consists of two distinct components: a rapid inward current carried by sodium ions and a slower activating outward current carried by potassium ions. These currents result from independent permeability mechanisms for Na^+^ and K^+^, where the conductance changes over time and membrane potential. Consequently, the model can replicate and explain a wide range of phenomena, including the shape and propagation of action potentials, the sharp threshold, refractory period, anode-break excitation, accommodation, and subthreshold oscillations. Minor adjustments in key conductance and stimulus current parameters enable the model to describe various action potential phenomena (*^52^*). The formula shows as follows:

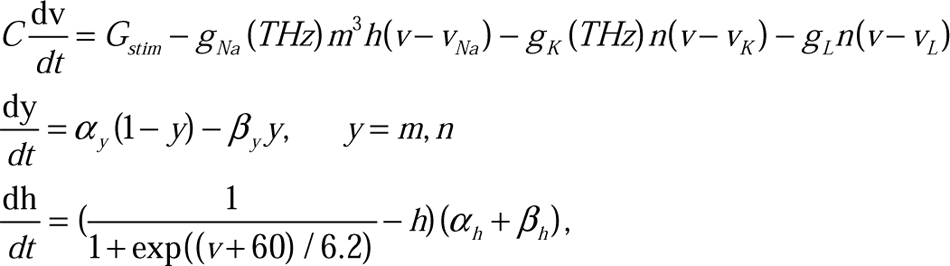

Where v, m, h and n represent the membrane voltage, probability of opening or closing of potassium-sodium ion channel. *V_Na_*, *V_K_* and *V_L_* are the sodium ion reverse potential, potassium ion reverse potential and resting membrane potential, respectively. *g_Na_*, *g_K_*are the maximum conductivity of sodium and potassium ion, respectively. C is membrane capacitive reactance with 0.75 uF/cm^2^, the *G_stim_* is the stimulation by an external current.

### High Frequency Terahertz and Blue Light Stimulation

For high frequency of terahertz stimulation (HFTS), we used a quantum cascade laser with a center frequency of 35.93±0.1 THz. The laser beam was coupled into a coupler, supported by the Innovation Laboratory of Terahertz Biophysics. We then connected the coupler to a Polycrystalline fiber (PIR) infrared fiber (Art photonics) with a core composition of AgCl/Br. This fiber has excellent transmittance in the range of 3-18 μm, with a core refractive index of 2.15 and an effective numerical aperture (NA) of 0.35 ± 0.05. At the distal end of the fiber, we left approximately 3-4 cm of bare fiber to allow for the insertion of a hollow tube with an inner diameter of 650 μm. This tube was pre-implanted into the ACC region of the SNI and sham group mice brain (*^53^*). The duration of HFTS was 15 minutes, with a pulse width of 2 μs, a repetition frequency of 10 kHz, and a duty cycle of 40%. The average output power at the tip of the fiber, measured by a MIR detector (NOVA II-3A, Israel), was 0.3 ± 0.05 mW. For comparison purposes, we also used a blue laser to stimulate the same brain region for 15 minutes, with a frequency of 1 Hz and an average output power of 10 mW.

### *In Vitro* Patch Clamp Recording

The experimental procedures were based on our previous reports (*^54^*). Briefly, mice were anesthetized and then decapitated to sacrifice. Brain slices (300 μm thick) containing the ACC were cut on a vibrating microtome (Leica VT 1200s, Heidelberger, Nussloch, Germany) at 0−4°C in oxygenated (95% O_2_ and 5% CO_2_) artificial cerebrospinal fluid (ACSF) consisting of (in mM) 124 NaCl, 25 NaHCO_3_, 2.5 KCl, 1 NaH_2_PO_4_, 2 CaCl_2_, 1 MgSO_4_ and 10 glucose. Slices were then transferred to a room temperature-submerged recovery chamber containing oxygenated ACSF and incubated for at least one hour before patch clamp recording. The neurons were then visualized under a microscope with infrared differential interference contrast or fluorescent optics video microscopy (BX51W1, Olympus, Tokyo, Japan). The recording pipettes (3-5 MΩ) were filled with a solution composed of (in mM) 124 K-gluconate, 5 NaCl, 1 MgCl_2_, 0.2 EGTA, 2 MgATP, 0.1 Na_3_GTP, 10 HEPES and 10 phosphocreatine disodium (adjusted to pH 7.2 with KOH, 290 mOsmol). Biocytin (0.2%) were added into pippette solution for verifying neurons and visualized through biocytin-avidin reaction. To examine the properties of voltage-gated K^+^ currents, tetrodotoxin (TTX, 1 μM) and CdCl_2_ (100 μM) were added into the ACSF. BaCl_2_ (4 mM) were applicated for recording K_Leak_ currents, since barium blocks most K_Leak_ channel subtypes. To examine voltage-gated Na^+^ current, 3 mM 4-AP and 0.1 mM CdCl_2_ were added into the ACSF. Electrical signals were filtered at 1 kHz by a Multiclamp 700B amplifier (Molecular Devices, USA), and digitized by an Axon DigiData 1550A converter with a sampling frequency of 10 kHz. Data analyses were performed with the Clampfit 10.02.

### *In Vivo* Multi-Channel Recording

Before the SNI operation, we implanted an electrode into the right ACC as in our previous reports (*^19^*), following stereotaxic coordinates: 1.1 mm anterior to the bregma, 0.3 mm lateral to the midline and 1.8 mm vertical to the skull surface. The electrodes were secured to the exposed skull using the dental adhesive resin cement Super-bond C&B (Japan)(*^53^*). This electrode consisted of 16-channel wire electrodes and included a hollow tube. During the optical stimulation, we employed multi-channel recording technology by Neurolego system (Nanjing Greathink Medical Technology, Nanjing, China). Subsequently, single-unit spike sorting was performed using the MClust-v4.4 toolbox in MATLAB software (MathWorks, USA). In the ACC region, the two main cell types are pyramidal neurons and interneuron cells, which are gamma-aminobutyric acid (GABA) neurons. Pyramidal neurons were primarily classified based on a trough-to-peak duration above 430 us, indicating long-duration action potentials. Interneuron cells, on the other hand, were identified based on a duration time below 430 μs (*^55^*).

### Behavioral Assays Mechanical allodynia

Briefly, the paw withdrawal mechanical threshold (PWMT) was evaluated by using von Frey filaments (Stoelting, Kiel, WI, USA) as reported in our previous works (*^54^*). Mice were habituated to the testing environment for 3 days before baseline testing and then placed under inverted plastic boxes (7 × 7 × 10 cm) on an elevated mesh floor and allowed to habituate for 30 min before threshold testing. A logarithmic series of 8 calibrated Semmes-Weinstein monofilaments (von Frey hairs; Stoelting, Kiel, WI, USA) (0.008, 0.02, 0.04, 0.16, 0.4, 0.6, 1, 1.4, and 2 g) with various bending forces (0.078, 0.196, 0.392, 1.568, 3.92, 5.88, 9.8, 13.72, and 19.6 mN) was applied to the plantar surface of the hind paw until the mice withdrew from the stimulus. Positive responses included licking, biting, and sudden withdrawal of the hind paws. A von Frey filament was applied 5 times (3 seconds for each stimulus) to each tested area. The minimum bending force of the von Frey filament able to evoke 3 occurrences of the paw withdrawal reflex was considered the paw withdrawal threshold. All tests were performed in a blinded manner.

### CatWalk gait analysis

Gait analysis was conducted using the CatWalk XT system (Noldus, the Netherlands) to measure pain-related parameters. The experimental setup involved placing the mouse on a glass platform with open ends, allowing the mouse to walk voluntarily. Simultaneously, a high-speed camera positioned underneath the platform captured images of each step, which were then transmitted to the analysis software (version 10.6, CatWalk XT, Noldus) for further processing. In this study, eight parameters were identified to assess dynamic behaviors relevant to neuropathic pain: (1) Stand: this parameter represents the duration (in seconds) of a paw touching the glass plate; (2) Stand index: it describes the speed at which the paw moves away from the glass plate; (3) Max contact area: it describes the maximum contact area of the paw or leg with the glass plate; (4) Mean print area: it represents the average area of the paw print during locomotion; (5) Mean intensity: this parameter denotes the average intensity value of the running stage; (6) Duty cycle This parameter denotes the average intensity value of the running stage.

### Statistical Analysis

GraphPad Prism 5 (Graph Pad Software, Inc.) was used for the statistical analyses and graphing. Statistical significance was assessed by unpaired *t-*test, paired *t*-test, one-way and two-way ANOVA followed by *post hoc* comparison, Wilcoxon matched-paired signed rank test and Mann-Whitney test. All data in the experiment are expressed in mean ± S.E.M. Statistical significance was indicated as *, *P* < 0.05, **, *P* < 0.01, ***, *P* < 0.001 and ****, *P* < 0.0001.

## Author contributions

W.-Y.P., Data curtation, Formal analysis, Investigation, Visualization, Writing-original draft, Writing-review and editing, Funding acquisition; P.W., Data curation, Investigation, Methodology; C.-Y.T., Data curation, Investigation, Methodology, Writing-original draft; H.Z., Investigation, Methodology; K.C., Investigation; H.-X.S., Methodology; Y.-C.T., Methodology; A.-X.L., Methodology; Z.Z., Investigation, software; Y.-F.Y., Investigation; K.-J.W., Data curation, Investigation, Methodology, Writing-original draft; C.C., Conceptualization, Supervision; Y.-M.W., Supervision, Investigation, Methodology; T.C., Conceptualization, Supervision, Funding acquisition, Investigation, Visualization, Methodology, Writing-review and editing, Project administration, Funding acquisition.

## Acknowledgments

We thank the support the teaching center of Air force medical university for the invaluable technical assistance and J.-X.G. of our laboratory for the helpful comments on an earlier version of the manuscript and discussions. This work was supported by grants from the National Natural Science Foundation of China (32192410, 32071000 to T. C., 82271893 to Y.-M.W., 8230071226 to W.-Y.P., 82302090 to K.C.), National Science Fund for Distinguished Young Scholars (12225511 to C.C.), National Science Fund of China Major Project (T2241002 to C.C.), Innovation Laboratory of Terahertz Biophysics (23-163-00-GZ-001-001-02-04 to W.-Y.P.), The Key Research and Development Plan of Shaanxi Province (S2024-YF-YBSF-0277 to W.-Y.P.)

## References

1. R. Baron, G. Binder A Fau - Wasner, G. Wasner, Neuropathic pain: diagnosis, pathophysiological mechanisms, and treatment. The Lancet. Neurology 9, 807–819 (2010).

2. L. Shan, D. Selvarajah, Z. Wang, Editorial: Insights in neuropathic pain: 2022. Frontiers in pain research (Lausanne, Switzerland) 4, 1232025 (2023).

3. T. S. Jensen, N. B. Finnerup, Allodynia and hyperalgesia in neuropathic pain: clinical manifestations and mechanisms. The Lancet. Neurology 13, 924–935 (2014).

4. M. Costigan, C. J. Scholz J Fau - Woolf, C. J. Woolf, Neuropathic pain: a maladaptive response of the nervous system to damage. Annual review of neuroscience 32, 1–32 (2009).

5. K. Liu, L. Wang, Optogenetics: Therapeutic spark in neuropathic pain. Bosnian journal of basic medical sciences 19, 321–327 (2019).

6. W. Peng et al., High-frequency terahertz waves disrupt Alzheimer’s β-amyloid fibril formation. eLight 3, 18 (2023).

7. Y. Li, C. Chang, Z. Zhu, & C. Fan, Terahertz Wave Enhances Permeability of the Voltage-Gated Calcium Channel. Journal of the American Chemical Society 143, 4311–4318 (2021).

8. Y. Zhao, L. Wang, Y. Li, Z. Zhu, Terahertz Waves Enhance the Permeability of Sodium Channels. Symmetry 15, 427 (2023).

9. J. Zhang et al., Non-invasive, opsin-free mid-infrared modulation activates cortical neurons and accelerates associative learning. Nature Communications 12, 2730 (2021).

10. X. Liu, Z. Qiao, Y. Chai, Z. Zhu, et al., Nonthermal and reversible control of neuronal signaling and behavior by midinfrared stimulation. Proceedings of the National Academy of Sciences of the United States of America 118, 10 (2021).

11. J. S. Trimmer, Ion channels and pain: important steps towards validating a new therapeutic target for neuropathic pain. Experimental neurology 254, 190–194 (2014).

12. T. V. P. Bliss, G. L. Collingridge, B.-K. Kaang, M. Zhuo, Synaptic plasticity in the anterior cingulate cortex in acute and chronic pain. Nature Reviews Neuroscience 17, 485–496 (2016).

13. M. A.-O. Tsuda, K. Koga, T. Chen, M. Zhuo, Neuronal and microglial mechanisms for neuropathic pain in the spinal dorsal horn and anterior cingulate cortex. Journal of neurochemistry 141(4), 486–498 (2017).

14. W. Chen T Fau - Wang et al., Postsynaptic insertion of AMPA receptor onto cortical pyramidal neurons in the anterior cingulate cortex after peripheral nerve injury. Molecular brain 7, 76 (2014).

15. K. Koga et al., Coexistence of two forms of LTP in ACC provides a synaptic mechanism for the interactions between anxiety and chronic pain. Neuron 85(2), 377–389 (2015).

16. T. Chen et al., Top-down descending facilitation of spinal sensory excitatory transmission from the anterior cingulate cortex. Nature communications 9(1), 1886 (2018).

17. S. R. A. Alles, P. A. Smith, Etiology and Pharmacology of Neuropathic Pain. Pharmacological reviews 70(2), 315–347 (2018).

18. T. V. Bliss, G. L. Collingridge, B. K. Kaang, M. Zhuo, Synaptic plasticity in the anterior cingulate cortex in acute and chronic pain. Nature reviews. Neuroscience 17 (8), 485–496 (2016).

19. D. Y. Zhu et al., The increased in vivo firing of pyramidal cells but not interneurons in the anterior cingulate cortex after neuropathic pain. Molecular brain 15(1), 12 (2022).

20. X.-Y. Li et al., Alleviating Neuropathic Pain Hypersensitivity by Inhibiting PKMζ in the Anterior Cingulate Cortex. Science 330, 1400–1404 (2010).

21. S. J. Kang et al., Bidirectional modulation of hyperalgesia via the specific control of excitatory and inhibitory neuronal activity in the ACC. Molecular Brain 8, 81 (2015).

22. P. N. Fuchs, Y. B. Peng, J. A. Boyette-Davis, M. L. Uhelski, The anterior cingulate cortex and pain processing. Frontiers in integrative neuroscience 8, 35 (2014).

23. K. Lançon, C. Qu, E. Navratilova, F. Porreca, P. Séguéla, Decreased dopaminergic inhibition of pyramidal neurons in anterior cingulate cortex maintains chronic neuropathic pain. Cell reports 37(9), 109933 (2021).

24. S. W. Um, M. J. Kim, J. W. Leem, S. J. Bai, B. H. Lee, Pain-Relieving Effects of mTOR Inhibitor in the Anterior Cingulate Cortex of Neuropathic Rats. Molecular Neurobiology 56, 2482–2494 (2019).

25. W. Tan et al., Alleviating neuropathic pain mechanical allodynia by increasing Cdh1 in the anterior cingulate cortex. Molecular pain 11, 56 (2015).

26. M. Takeda et al., Potassium channels as a potential therapeutic target for trigeminal neuropathic and inflammatory pain. Molecular pain 7, 5 (2011).

27. X. Du, N. Gamper, Potassium channels in peripheral pain pathways: expression, function and therapeutic potential. Current neuropharmacology 11(6), 621–40 (2013).

28. A. Sakai et al., MicroRNA cluster miR-17-92 regulates multiple functionally related voltage-gated potassium channels in chronic neuropathic pain. Nature Communications 8, 16079 (2017).

29. J.-Y. Zhao et al., DNA methyltransferase DNMT3a contributes to neuropathic pain by repressing Kcna2 in primary afferent neurons. Nature Communications 8, 14712 (2017).

30. L. Fan et al., Impaired neuropathic pain and preserved acute pain in rats overexpressing voltage-gated potassium channel subunit Kv1.2 in primary afferent neurons. Molecular Pain 10, 8 (2014).

31. C. Tsantoulas, S. B. McMahon, Opening paths to novel analgesics: the role of potassium channels in chronic pain. Trends in neurosciences 37(3), 146–158 (2014).

32. N. A.-O. Khandelwal et al., FOXP1 negatively regulates intrinsic excitability in D2 striatal projection neurons by promoting inwardly rectifying and leak potassium currents. Molecular psychiatry 26(6), 1761–1774 (2021).

33. C. González et al., K(+) channels: function-structural overview. Comprehensive Physiology 2(3), 2087–2149 (2012).

34. K. L. Zhang et al., Targeted up-regulation of Drp1 in dorsal horn attenuates neuropathic pain hypersensitivity by increasing mitochondrial fission. Redox biology 49, 102216 (2022).

35. R. Zhao et al., Neuropathic Pain Causes Pyramidal Neuronal Hyperactivity in the Anterior Cingulate Cortex. Frontiers in cellular neuroscience 12, 107 (2018).

36. Z. Chen, X. Shen, L. Huang, H. Wu, M. Zhang, Membrane potential synchrony of neurons in anterior cingulate cortex plays a pivotal role in generation of neuropathic pain. Scientific Reports 8, 1691 (2018).

37. H. Xu et al., Presynaptic and postsynaptic amplifications of neuropathic pain in the anterior cingulate cortex. The Journal of neuroscience: the official journal of the Society for Neuroscience 28(29), 7445–7453 (2008).

38. K. Gao, Z. Lin, S. Wen, Y. Jiang, Potassium channels and epilepsy. Acta neurologica Scandinavica 146(6), 699–707 (2022).

39. S. Maljevic, H. Lerche, Potassium channels: a review of broadening therapeutic possibilities for neurological diseases. Journal of neurology 260(9), 2201–2211 (2013).

40. A. Al-Sabi, Seshu K. Kaza, J. O. Dolly, J. Wang, Pharmacological characteristics of Kv1.1- and Kv1.2-containing channels are influenced by the stoichiometry and positioning of their α subunits. Biochemical Journal 454, 101–108 (2013).

41. Y. Liang et al., Transcription factor EBF1 mitigates neuropathic pain by rescuing Kv1.2 expression in primary sensory neurons. Translational research: the journal of laboratory and clinical medicine 263, 15–27 (2024).

42. J. Zhang et al., Epigenetic restoration of voltage-gated potassium channel Kv1.2 alleviates nerve injury-induced neuropathic pain. Journal of neurochemistry 156, 367–378 (2021).

43. T. A.-O. Xiao et al., Sensory input-dependent gain modulation of the optokinetic nystagmus by mid-infrared stimulation in pigeons. eLife 12, e78729 (2023).

44. X. Xiao, Y.-Q. Zhang, A new perspective on the anterior cingulate cortex and affective pain. Neuroscience & Biobehavioral Reviews 90, 200–211 (2018).

45. S. J. Kang et al., Bidirectional modulation of hyperalgesia via the specific control of excitatory and inhibitory neuronal activity in the ACC. Molecular brain 8(1), 81 (2015).

46. J. Cichon, T. J. J. Blanck, W.-B. Gan, G. Yang, Activation of cortical somatostatin interneurons prevents the development of neuropathic pain. Nature Neuroscience 20, 1122–1132 (2017).

47. R. Zhou, W. Wang C Fau - Xu, L. Xu W Fau - Xie, L. Xie, Biological applications of terahertz technology based on nanomaterials and nanostructures. Nanoscale 11(8), 3445–3457 (2019).

48. Q. Wang, Y. Chen, J. Mao, F. Yang, N. A.-O. Wang, Metasurface-Assisted Terahertz Sensing. Sensors (Basel, Switzerland) 23(13), 5902 (2023).

49. B. A. Cole, S. J. Clapcote, S. P. Muench, J. D. Lippiat, Targeting K(Na)1.1 channels in KCNT1-associated epilepsy. Trends in pharmacological sciences 42(8), 700–713 (2021).

50. T. J. Jentsch, Neuronal KCNQ potassium channels: physiology and role in disease. Neuroscience 1(1), 21–30 (2000).

51. J. L. Taylor et al., Functionally linked potassium channel activity in cerebral endothelial and smooth muscle cells is compromised in Alzheimer’s disease. Proceedings of the National Academy of Sciences of the United States of America 119(26), (2022).

52. M. Häusser, The Hodgkin-Huxley theory of the action potential. Nature Neuroscience 3, 1165–1165 (2000).

53. Z. V. Guo et al., Procedures for behavioral experiments in head-fixed mice. PloS one 9(2), e101397 (2014).

54. M. M. Zhang et al., Glutamatergic synapses from the insular cortex to the basolateral amygdala encode observational pain. Neuron 110,1993–2008 (2022).

55. A. V. Apkarian, R.-D. Bushnell Mc Fau - Treede, J.-K. Treede Rd Fau - Zubieta, J. K. Zubieta, Human brain mechanisms of pain perception and regulation in health and disease. European journal of pain (London, England) 9(4), 463–484 (2005).

