## Supplement data and tables for "High Frequency Terahertz Stimulation Alleviates Neuropathic Pain by Inhibiting the Pyramidal Neuron Activity in the Anterior Cingulate Cortex of mice"

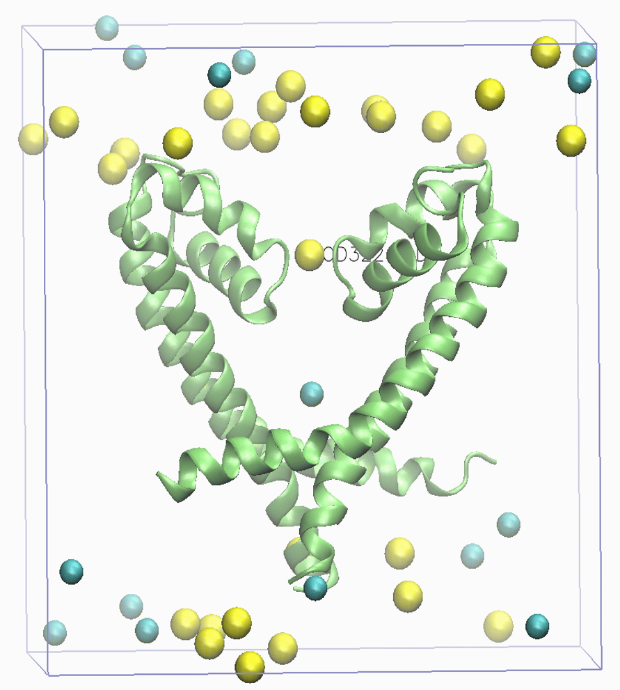

Figure S1 Sodium (Na^+^) channels (PDB ID: 3RVY) developed in this study with clean view. Yellow balls represent the Na^+^, blue balls represent the Cl^-^.

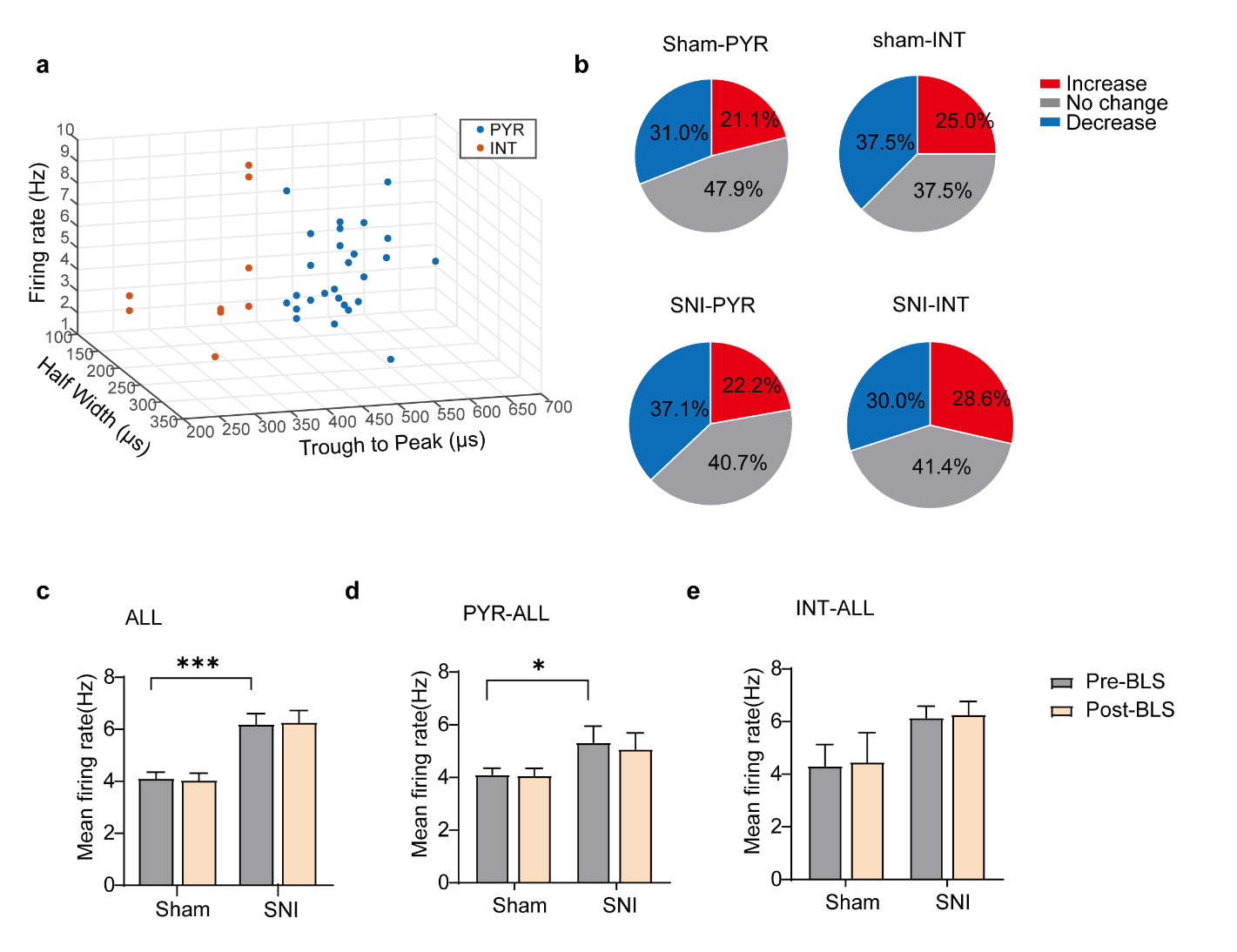

Figure S2 Blue light stimulation (BLS) has no effect on the mean firing rate on ACC neurons in both sham and SNI mice. (a) Partial recorded ACC neurons were classified as pyramidal (PYR) cells and interneurons (INT) using k-means cluster-separation algorithm based on their electrophysiological properties. (b) Pie charts summarize the changes in firing rate of PYR (70/78, 27/97) and INT (*n* = 8/78, 70/97) in sham (*n* = 78) and SNI (*n* = 97) groups. Pre vs. post BLS, Wilcoxon rank-sum test and paired *t*-test. (c) The mean firing rate of all neurons in sham and SNI groups before and after BLS. Sham group (*P* >0.05, Wilcoxon matched-paired signed rank test), SNI group (*P* > 0.05, Wilcoxon matched-paired signed rank test), SNI pre-HFTS vs. Sham pre-HFTS (*P* < 0.001, Mann-Whitney test). (d) The mean firing rate of PYR neurons in sham and SNI groups before and after BLS. Sham group (*P* > 0.05, Wilcoxon matched-paired signed rank test), SNI group (*P* > 0.05, Wilcoxon matched-paired signed rank test), SNI pre-HFTS vs. Sham pre-HFTS (*P* < 0.05, Mann-Whitney test). (e)The mean firing rate of INT neurons in sham and SNI groups before and after BLS. Sham group (*P* >0.05, Wilcoxon matched-paired signed rank test), SNI group (*P* > 0.05, Wilcoxon matched-paired signed rank test). SNI pre-HFTS vs. Sham pre-HFTS (*P* > 0.05, Mann-Whitney test).*, *P* < 0.05, ****, *P* < 0.0001, ns, *P* > 0.05. PYR, pyramidal neurons. SNI, spared nerve injury.

| Table S1-1. The Kv current to different voltage in PYR of SNI mice before and after HFTS | | | | | |
| --- | --- | --- | --- | --- | --- |
| SNI - SNI+HFTS | Mean Diff. | 95.00% CI of diff. | Below threshold? | Summary | Adjusted P Value |
| -110 mV | 35.81 | -1266 to 1338 | No | ns | >0.9999 |
| -100 mV | 91.01 | -1211 to 1393 | No | ns | >0.9999 |
| -90 mV | 51.45 | -1250 to 1353 | No | ns | >0.9999 |
| -80 mV | -1.271 | -1303 to 1301 | No | ns | >0.9999 |
| -70 mV | 37.7 | -1264 to 1339 | No | ns | >0.9999 |
| -60 mV | 21.7 | -1280 to 1323 | No | ns | >0.9999 |
| -50 mV | 35.37 | -1266 to 1337 | No | ns | >0.9999 |
| -40 mV | 58.42 | -1243 to 1360 | No | ns | >0.9999 |
| -30 mV | 159.3 | -1142 to 1461 | No | ns | >0.9999 |
| -20 mV | 208.7 | -1093 to 1510 | No | ns | >0.9999 |
| -10 mV | -25.13 | -1327 to 1277 | No | ns | >0.9999 |
| 0 mV | -98.64 | -1400 to 1203 | No | ns | >0.9999 |
| 10 mV | -529.7 | -1832 to 772.0 | No | ns | 0.9969 |
| 20 mV | -830.9 | -2133 to 470.9 | No | ns | 0.7063 |
| 30 mV | -1010 | -2311 to 292.1 | No | ns | 0.3384 |
| 40 mV | -1294 | -2596 to 7.778 | No | ns | 0.053 |
| 50 mV | -1471 | -2773 to -169.7 | Yes | * | 0.0127 |
| 60 mV | -1726 | -3028 to -424.5 | Yes | ** | 0.0012 |
| 70 mV | -1816 | -3118 to -514.3 | Yes | *** | 0.0005 |
| 80 mV | -1856 | -3157 to -553.8 | Yes | *** | 0.0003 |
| 90 mV | -1900 | -3202 to -598.7 | Yes | *** | 0.0002 |
| 100 mV | -1839 | -3141 to -537.0 | Yes | *** | 0.0004 |
| 110 mV | -1915 | -3216 to -612.7 | Yes | *** | 0.0002 |
| 120 mV | -1912 | -3214 to -610.4 | Yes | *** | 0.0002 |
| 130 mV | -1912 | -3214 to -610.0 | Yes | *** | 0.0002 |
| ANOVA table | SS | DF | MS | F (DFn, DFd) | P value |
| Row Factor x Column Factor | 55521178 | 24 | 2313382 | F (24, 240) = 8.581 | P<0.0001 |
| Row Factor | 511369305 | 24 | 21307054 | F (24, 240) = 79.03 | P<0.0001 |
| Column Factor | 45335332 | 1 | 45335332 | F (1, 10) = 6.846 | P=0.0258 |
| Subject | 66223673 | 10 | 6622367 | F (10, 240) = 24.56 | P<0.0001 |
| Residual | 64706213 | 240 | 269609 |  |  |
| Data summary | Šídák's multiple comparisons test | | | | |
| Number of columns (Column Factor) | 2 | | | | |
| Number of rows (Row Factor) | 25 | | | | |
| Number of subjects (Subject) | 12 | | | | |
| Number of missing values | 0 | | | | |

| Table S1-2. Changes of the Kv current impact by HFTS over time | | | | | |
| --- | --- | --- | --- | --- | --- |
|  | Mean Diff. | 95.00% CI of diff. | Below threshold? | Summary | Adjusted P Value |
| Pre HFTS vs. 5 min | -899.9 | -1611 to -188.4 | Yes | * | 0.0103 |
| Pre HFTS vs. 20 min | -790.3 | -1502 to -78.79 | Yes | * | 0.026 |
| 5 min vs. 20min | 109.6 | -601.9 to 821.1 | No | ns | 0.9953 |
| ANOVA table | SS | DF | MS | F (DFn, DFd) | P value |
| Treatment (between columns) | 1969090 | 4 | 492273 | F (4, 15) = 4.193 | P=0.0178 |
| Residual (within columns) | 1761043 | 15 | 117403 |  |  |
| Total | 3730134 | 19 |  |  |  |
| Data summary | Šídák's multiple comparisons test | | | | |
| Number of treatments (columns) | 5 | | | | |
| Number of values (total) | 20 | | | | |

| Table S1-3. The K_leak_ current to different voltage in PYR of SNI mice before and after HFTS | | | | | |
| --- | --- | --- | --- | --- | --- |
| SNI - SNI+HFTS | Mean Diff. | 95.00% CI of diff. | Below threshold? | Summary | Adjusted P Value |
| -120 mV | 19.22 | -2.693 to 41.14 | No | ns | 0.129 |
| -110 mV | 14.26 | -7.660 to 36.17 | No | ns | 0.4949 |
| -100 mV | 11.54 | -10.37 to 33.46 | No | ns | 0.7675 |
| -90 mV | 6.985 | -14.93 to 28.90 | No | ns | 0.9894 |
| -80 mV | 5.231 | -16.68 to 27.15 | No | ns | 0.999 |
| -70 mV | -1.408 | -23.32 to 20.51 | No | ns | >0.9999 |
| -60 mV | -7.848 | -29.76 to 14.07 | No | ns | 0.9752 |
| -50 mV | -14.48 | -36.40 to 7.432 | No | ns | 0.472 |
| -40 mV | -30.52 | -52.43 to -8.600 | Yes | ** | 0.0012 |
| -30 mV | -46.87 | -68.79 to -24.95 | Yes | **** | <0.0001 |
| ANOVA table | SS | DF | MS | F (DFn, DFd) | P value |
| Row Factor x Column Factor | 13968 | 9 | 1552 | F (9, 108) = 8.377 | P<0.0001 |
| Row Factor | 122682 | 9 | 13631 | F (9, 108) = 73.58 | P<0.0001 |
| Column Factor | 674.2 | 1 | 674.2 | F (1, 12) = 1.688 | P=0.2182 |
| Subject | 4792 | 12 | 399.3 | F (12, 108) = 2.156 | P=0.0189 |
| Residual | 20008 | 108 | 185.3 |  |  |
| Data summary | Šídák's multiple comparisons test | | | | |
| Number of columns (Column Factor) | 2 | | | | |
| Number of rows (Row Factor) | 10 | | | | |
| Number of subjects (Subject) | 14 | | | | |
| Number of missing values | 0 | | | | |

| Table S2-1. The spike frequency in PYR of ACC before and after SNI | | | | | |
| --- | --- | --- | --- | --- | --- |
| Table Analyzed | Sham vs SNI | | | | |
| Two-way RM ANOVA | Matching: Across row | | | | |
| Assume sphericity? | Yes | | | | |
| Alpha | 0.05 | | | | |
| Source of Variation | % of total variation | P value | P value summary | Significant? |  |
| Row Factor x Column Factor | 1.652 | 0.1771 | ns | No |  |
| **Row Factor** | **54.75** | **<0.0001** | ******** | **Yes** |  |
| **Column Factor** | **18.86** | **<0.0001** | ******** | **Yes** |  |
| Subject | 18.67 | 0.0003 | *** | Yes |  |
| ANOVA table | SS | DF | MS | F (DFn, DFd) | P value |
| Row Factor x Column Factor | 121.6 | 7 | 17.37 | F (7, 40) = 1.555 | P=0.1771 |
| Row Factor | 4030 | 7 | 575.7 | F (7, 40) = 16.76 | P<0.0001 |
| Column Factor | 1388 | 1 | 1388 | F (1, 40) = 124.2 | P<0.0001 |
| Subject | 1374 | 40 | 34.35 | F (40, 40) = 3.075 | P=0.0003 |
| Residual | 446.9 | 40 | 11.17 |  |  |
| Difference between column means |  | | | | |
| Mean of Con | 9.792 | | | | |
| Mean of SNI | 17.4 | | | | |
| Difference between means | -7.604 | | | | |
| SE of difference | 0.6823 | | | | |
| 95% CI of difference | -8.983 to -6.225 | | | | |
| Data summary |  | | | | |
| Number of columns (Column Factor) | 2 | | | | |
| Number of rows (Row Factor) | 8 | | | | |
| Number of subjects (Subject) | 48 | | | | |
| Number of missing values | 0 | | | | |

| Table S2-2. The spike frequency in PYR of SNI mice before and after HFTS | | | | | |
| --- | --- | --- | --- | --- | --- |
| Table Analyzed | SNI vs HFTS | | | | |
| Two-way RM ANOVA | Matching: Across row | | | | |
| Assume sphericity? | Yes | | | | |
| Alpha | 0.05 | | | | |
| Source of Variation | % of total variation | P value | P value summary | Significant? |  |
| Row Factor x Column Factor | 0.461 | 0.995 | ns | No |  |
| **Row Factor** | **36.3** | **<0.0001** | ******** | **Yes** |  |
| **Column Factor** | **11.29** | **<0.0001** | ******** | **Yes** |  |
| Subject | 32.41 | 0.0567 | ns | No |  |
| ANOVA table | SS | DF | MS | F (DFn, DFd) | P value |
| Row Factor x Time | 33.33 | 7 | 4.762 | F (7, 40) = 0.1349 | P=0.9950 |
| Row Factor | 2625 | 7 | 375 | F (7, 40) = 6.399 | P<0.0001 |
| Time | 816.7 | 1 | 816.7 | F (1, 40) = 23.13 | P<0.0001 |
| Subject | 2344 | 40 | 58.59 | F (40, 40) = 1.659 | P=0.0567 |
| Residual | 1413 | 40 | 35.31 |  |  |
| Difference between column means |  | | | | |
| Mean of SNI | 18.49 | | | | |
| Mean of SNI+THz | 12.66 | | | | |
| Difference between means | 5.833 | | | | |
| SE of difference | 1.213 | | | | |
| 95% CI of difference | 3.382 to 8.285 | | | | |
| Data summary |  | | | | |
| Number of columns (Time) | 2 | | | | |
| Number of rows (Row Factor) | 8 | | | | |
| Number of subjects (Subject) | 48 | | | | |
| Number of missing values | 0 | | | | |

| Table S2-3. The spike frequency in PYR of SNI mice before and after BLS | | | | | |
| --- | --- | --- | --- | --- | --- |
| Table Analyzed | SNI vs BLS | | | | |
| Two-way RM ANOVA | Matching: Across row | | | | |
| Assume sphericity? | Yes | | | | |
| Alpha | 0.05 | | | | |
| Source of Variation | % of total variation | P value | P value summary | Significant? |  |
| Row Factor x Column Factor | 1.416 | 0.9018 | ns | No |  |
| Row Factor | 11.29 | 0.0635 | ns | No |  |
| Column Factor | 0.07243 | 0.7101 | ns | No |  |
| Subject | 0.461 | 0.9716 | ns | No |  |
| ANOVA table | SS | DF | MS | F (DFn, DFd) | P value |
| Row Factor x Column Factor | 183.3 | 7 | 26.19 | F (7, 40) = 0.3915 | P=0.9018 |
| Row Factor | 4361 | 7 | 623 | F (7, 40) = 4.362 | P=0.0011 |
| Column Factor | 9.375 | 1 | 9.375 | F (1, 40) = 0.1401 | P=0.7101 |
| Subject | 5714 | 40 | 142.8 | F (40, 40) = 2.135 | P=0.0092 |
| Residual | 2676 | 40 | 66.9 |  |  |
| Difference between column means |  | | | | |
| Mean of SNI | 22.55 | | | | |
| Mean of SNI+Visible light | 21.93 | | | | |
| Difference between means | 0.625 | | | | |
| SE of difference | 1.67 | | | | |
| 95% CI of difference | -2.749 to 3.999 | | | | |
| Data summary |  | | | | |
| Number of columns (Column Factor) | 2 | | | | |
| Number of rows (Row Factor) | 8 | | | | |
| Number of subjects (Subject) | 48 | | | | |
| Number of missing values | 0 | | | | |

| Table S2-4. The spike frequency in PYR of Sham mice before and after HFTS | | | | | |
| --- | --- | --- | --- | --- | --- |
| Table Analyzed | Sham vs HFTS | | | | |
| Two-way RM ANOVA | Matching: Across row | | | | |
| Assume sphericity? | Yes | | | | |
| Alpha | 0.05 | | | | |
| Source of Variation | % of total variation | P value | P value summary | Significant? |  |
| Row Factor x Column Factor | 4.831 | 0.0029 | ** | Yes |  |
| **Row Factor** | **66.5** | **<0.0001** | ******** | **Yes** |  |
| **Column Factor** | **15.75** | **<0.0001** | ******** | **Yes** |  |
| Subject | 5.691 | 0.7722 | ns | No |  |
| ANOVA table | SS | DF | MS | F (DFn, DFd) | P value |
| Row Factor x Column Factor | 342.6 | 7 | 48.95 | F (7, 40) = 3.824 | P=0.0029 |
| Row Factor | 4717 | 7 | 673.8 | F (7, 40) = 66.77 | P<0.0001 |
| Column Factor | 1117 | 1 | 1117 | F (1, 40) = 87.29 | P<0.0001 |
| Subject | 403.6 | 40 | 10.09 | F (40, 40) = 0.7884 | P=0.7722 |
| Residual | 512 | 40 | 12.8 |  |  |
| Difference between column means |  | | | | |
| Mean of Sham | 12.45 | | | | |
| Mean of Sham+HFST | 5.625 | | | | |
| Difference between means | 6.823 | | | | |
| SE of difference | 0.7303 | | | | |
| 95% CI of difference | 5.347 to 8.299 | | | | |
| Data summary |  | | | | |
| Number of columns (Column Factor) | 2 | | | | |
| Number of rows (Row Factor) | 8 | | | | |
| Number of subjects (Subject) | 48 | | | | |
| Number of missing values | 0 | | | | |

Note：PYR, pyramidal neurons; HFTS, high frequency terahertz stimulation; BLS, blue light stimulation
